# MicroRNA clusters integrate evolutionary constraints on expression and target affinities: the miR-6/5/4/286/3/309 cluster in *Drosophila* leg development

**DOI:** 10.1101/714766

**Authors:** Zhe Qu, Wing Chung Yiu, Ho Yin Yip, Wenyan Nong, Clare W.C. Yu, Ivy H.T. Lee, Annette Y.P. Wong, Nicola W.Y. Wong, Fiona K.M. Cheung, Ting Fung Chan, Kwok Fai Lau, Silin Zhong, Ka Hou Chu, Stephen S. Tobe, David E.K. Ferrier, William G. Bendena, Jerome H.L. Hui

## Abstract

A striking feature of microRNAs is that they are often clustered in the genomes of animals. The functional and evolutionary consequences of this clustering remain obscure. Here, we investigated a microRNA cluster miR-6/5/4/286/3/309 that is conserved across drosophilid lineages. Small RNA sequencing revealed expression of this microRNA cluster in *Drosophila melanogaster* leg discs, and conditional overexpression of the whole cluster resulted in leg appendage shortening. Transgenic overexpression lines expressing different combinations of microRNA cluster members were also constructed. Expression of individual microRNAs from the cluster resulted in a normal wild-type phenotype, but either the expression of several ancient microRNAs together (miR-5/4/286/3/309) or more recently evolved clustered microRNAs (miR-6-1/2/3) can recapitulate the phenotypes generated by the whole-cluster overexpression. Screening of transgenic fly lines revealed down-regulation of leg patterning gene cassettes in generation of the leg-shortening phenotype. Furthermore, cell transfection with different combinations of microRNA cluster members revealed a suite of downstream genes targeted by all cluster members, as well as complements of targets that are unique for distinct microRNAs. Considering together the microRNA targets and the evolutionary ages of each microRNA in the cluster demonstrates the importance of microRNA clustering, where new members can reinforce and modify the selection forces on both the cluster regulation and the gene regulatory network of existing microRNAs.

## Introduction

Operation of genes within multigenic clusters is widespread, but the functional and evolutionary implications of this are often poorly understood. In microbes, the poly-cistronic transcription of operons and anti-phage defensive system are well known for their importance (Doron et al 2018). In animals, there are various examples of protein-coding genes that are regulated within clusters. For example, homeobox genes in the Hox cluster are regulated through multigenic regulatory elements (Deschamps and Duboule 2017). In addition to these cases of clustered protein-coding genes, non-protein encoding genes such as those producing microRNAs are also often found to be in co-regulated clusters (e.g. Altuvia et al 2005; Fromm et al 2015; Lagos-Quintana et al 2001; Mohammed et al 2014; Bartel 2018). For instance, synchronised expression of clustered microRNAs in normal human cells is found to be mis-regulated during disease development (Dambal et al 2015; Nojima et al 2016), and mis-regulation has been implicated in cancer formation (Ventura et al 2008; Kim et al 2009). Since these microRNA clusters are relatively recent discoveries compared to protein-coding gene clusters, much less is known about the range of functional consequences of this clustering and the resultant evolutionary impacts.

There is an important difference between protein-coding versus microRNA gene clusters in animals. The individual genes in protein-coding gene clusters tend to have their own promoters, whereas microRNA clusters are often comprised of members transcribed as a single unit or polycistronic transcript regulated by a single promoter (Kozomara and Griffiths-Jones 2014). In addition, microRNA genes in a cluster are sometimes found to be conserved in sequence and orientation (e.g. the miR-17 cluster in mammals, Tanzer and Stadler 2004). This could be a consequence of *de novo* formation of hairpins in existing microRNA transcripts potentially being the major mechanism giving rise to new cluster members (Marco et al 2013; Wang et al 2016). Also, microRNAs in the same cluster are proposed to possess similar targeting properties or regulate genes in the same pathway (e.g. Kim and Nam 2006; Yuan et al 2009; Wang et al 2011; Hausser and Zavolan 2014; Wang et al 2016), although this remains somewhat controversial (e.g. Marco 2019; Wang et al 2019). In fact, the range of functional and evolutionary implications of polycistronic microRNAs in general remains controversial with regards to whether they are non-adaptive, the by-product of a tight genomic linkage, or simply expressed together due to unknown functional constraints (e.g. Marco et al 2013). A fundamental issue in these controversies is that most of these studies rely on correlating expression of the cluster with *in silico* prediction of their target genes. Systematic dissection of the functions of individual microRNAs from a cluster versus the function of the cluster as a whole remains to be tested.

## Results and Discussion

The miR-309-6 microRNA cluster has a distinctive composition/ organization of miR-6/5/4/286/3/309 in drosophilids that is conserved across the genus (Fig. 1A, Supplementary data S1). The miR-309-6 cluster contains different microRNA family members with different origins, such as miR-309 and miR-3 belong to MiR-3 family that originated in the Pancrustacea, miR-286 is in the Protostomia conserved MIR-279 family and miR-4 belongs to the most ancient MIR-9 family (Fromm et al 2019; Mohammed et al 2014; Ninova et al 2014). The cluster is located between genes CG15125 and CG11018, and is processed from a single ~1.5kb transcript in *Drosophila melanogaster* (Biemar et al 2005). It has high expression levels in early embryos (Ninova et al 2014) and deletion of the cluster results in partial larval lethality (Bushati et al 2008; Chen et al 2014). To identify the targets that are being regulated by miR-309-6 clusters in different drosophilids, *in silico* prediction of the microRNA targets of individual members of the cluster were carried out using miRanda and Targetscan (Supplementary data S2). In *D. melanogaster*, 37-38% of the total target genes were shared between at least 2 microRNAs in the cluster (Fig. 1B). In other drosophilid species, excluding the numbers predicted in *D. willistoni* because these are based on only a small number of available transcriptomes, 12.59%-22.77% of targets were shared between at least 2 microRNAs (Supplementary data S2). This is in agreement with previous data that suggested that microRNAs within a cluster may share common target genes (e.g. Kim and Nam 2006; Yuan et al 2009; Wang et al 2011; Hausser and Zavolan 2014; Wang et al 2016).

**Fig. 1.**
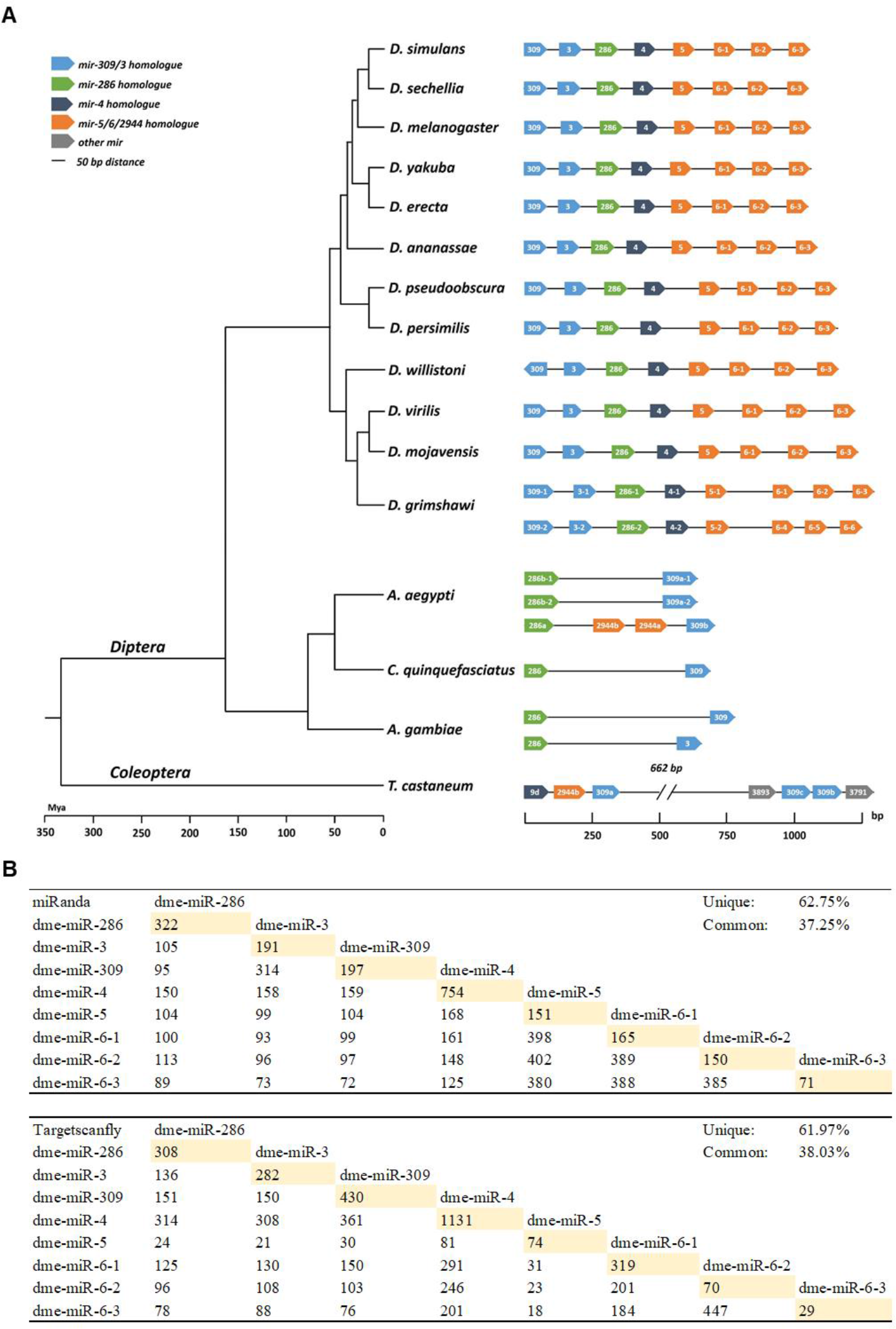
miR-309-6 cluster (miR-6/5/4/286/3/309) in various insects. A) Genomic organisation of the miR-309-6 cluster in insects (drawn to scale). MicroRNA homologues are shown by the same colour (Mohammed et al 2014; Ninova et al 2014, 2016). B) Number of miRanda and targetscanfly *in silico*-predicted target genes shared by microRNAs in the cluster in *Drosophila melanogaster*. Unique targets are highlighted in yellow. For other species refer to supplementary data S2.

As bioinformatic predictions of microRNA targets are prone to inclusion of false positives (e.g. Pinzon et al 2017), functional investigation was undertaken. Loss-of-function of the whole cluster was reported to result in larval lethality at different larval stages with about 57%-80% offspring survived to adulthood that were viable and fertile (Bushati et al 2008; Chen et al 2014), and in our hands this whole-cluster deletion line also resulted in partial larval lethality as previously reported and around 50% larvae survived to adults. Previous studies mainly focused on the functional importance of this microRNA cluster in embryonic stages (i.e. during the maternal-to-zygotic transition), and there is limited information on its function in other developmental stages or tissues (e.g. misorientation of adult sensory bristles on the adult notum with miR-3/-309 overexpression, Zhou et al 2018). We examined microRNA expression level/ pattern of this cluster in MirGeneDB and analysed the small RNA data sets of ModENCODE (Supplementary data S3), confirming that members of miR-309-6 cluster are expressing in various developmental stages/tissues. To facilitate further analyses in late development we sequenced the small RNA contents of leg discs in *D. melanogaster* L3 larvae, and revealed the expression of mature microRNAs contained in this cluster (Fig. 2). This finding was further validated by Taqman microRNA assays (Supplementary data S3), implying their potential functional roles during *D. melanogaster* leg development. The expression levels of miR-309 and miR-3 are lower than other members contained in the cluster, which is consistent with a recent study showing faster degradation rates of miR-309 and −3 (Zhou et al 2018).

**Fig. 2.**
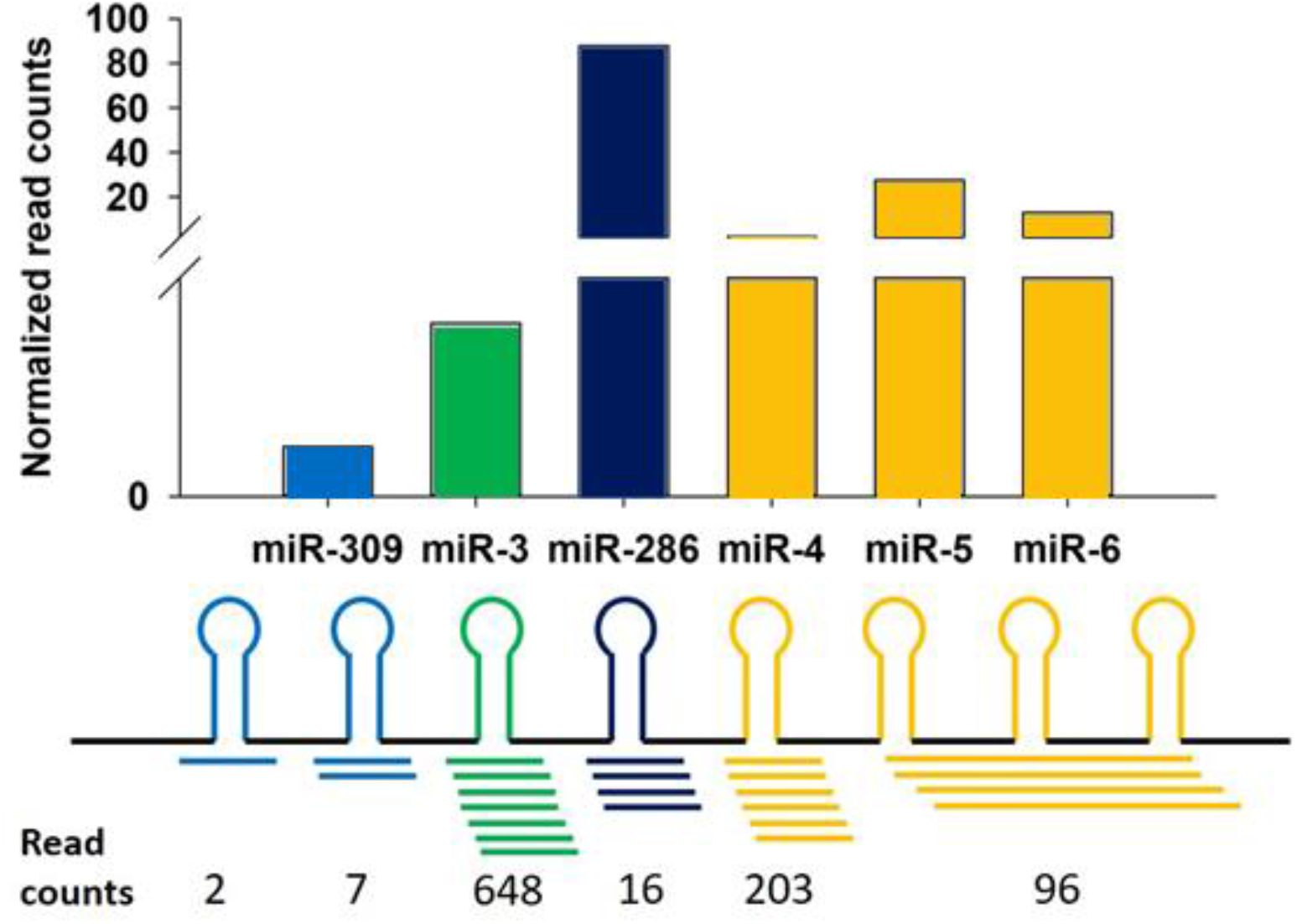
Small RNA sequencing revealed the expression of the miR-309-6 cluster in *D. melanogaster* L3 larvae leg discs. The clean read counts mapped by bowtie were shown (Upper panel: normalized read counts; lower panel: raw counts).

To enable functional analyses in late development we generated two homozygous UAS-miR-309-6 cluster lines, with the whole cluster independently inserted at 3L:3714826 (5’end of CG32264, named UAS-miR-309-6-I) and 3R:25235447 (5’end of CG10420, named UAS-miR-309-6-II). The location and orientation of these insertions was confirmed by Splinkerette PCR.

We screened for phenotypes by crossing UAS-miR-309-6-I and UAS-miR-309-6-II with GAL4 lines that targeted different tissues. Crossing UAS lines to either GAL4-*Dll* or GAL4-*ptc,* which targets *Distal-less* or *patched* expressing cells in the leg and wing discs, resulted in shortening and deletion of leg tarsal segments (Fig. 3B-C) relative to control animals (Fig. 3A-C, Supplementary information S4). Loss of the wing anterior cross-vein was also observed in these animals (Fig.3P-Q). Flies with shortened leg segments due to overexpression of the whole microRNA cluster were fertile and able to mate. Nevertheless, the mobility of both males and females was reduced. Moreover, during courtship, more effort was required for males and females to copulate, and the penetration time was dramatically reduced compared to wild type.

**Fig. 3.**
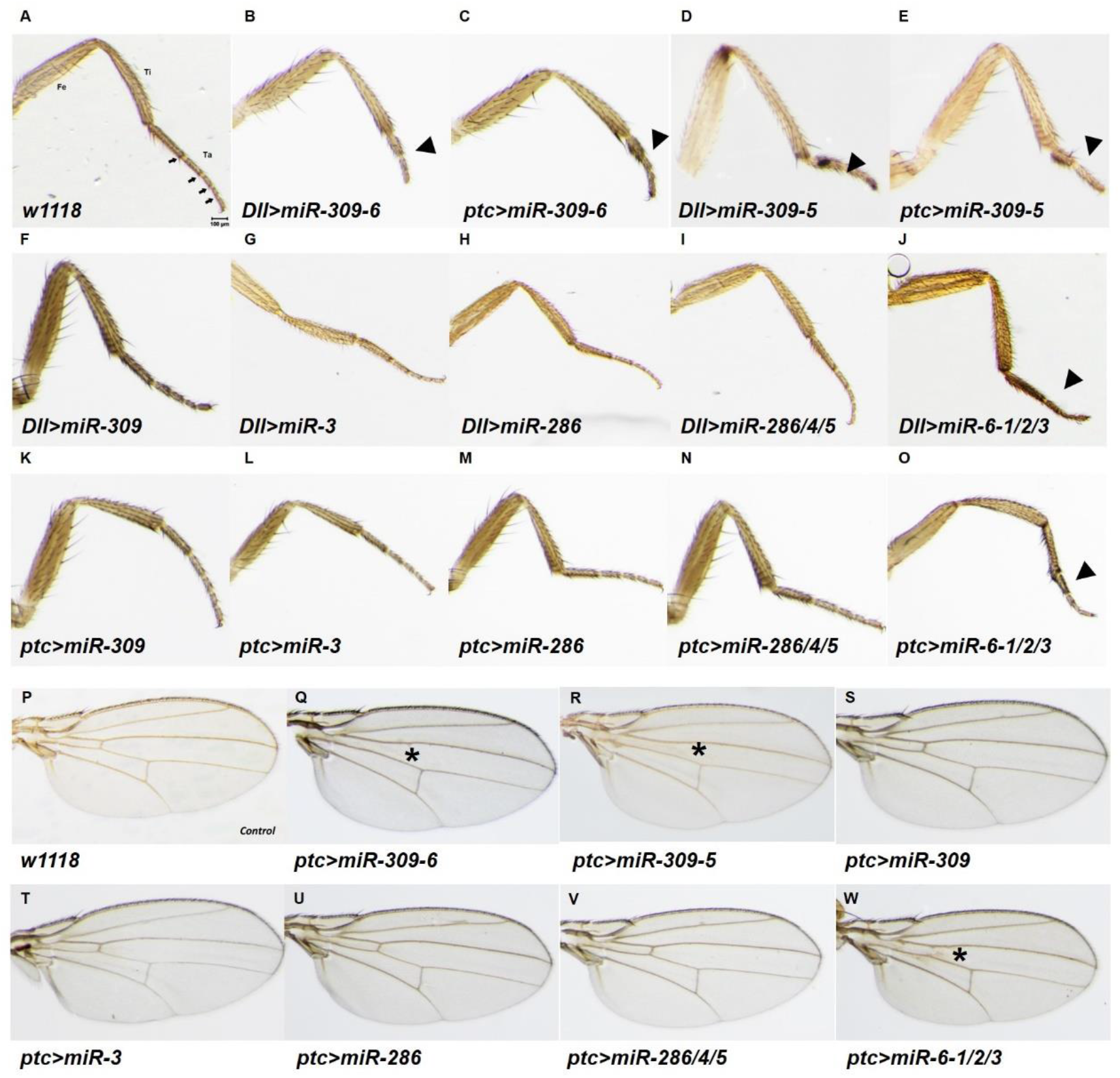
Expression of the whole miR-309-6 cluster, miR-309-5 or miR-6-1/2/3 sub-clusters results in shortening of leg tarsus and loss of anterior cross vein, whereas expression of other microRNAs from the cluster does not cause these phenotypic changes. Leg pictures of A) w1118 (control), B-E) expression of whole miR-309-6 cluster and miR-309-5 sub-cluster in either the *Distal-less* or *patched* expressing cells, F-O) expression of microRNAs from the cluster in either the *Distal-less* or *patched* expressing cells; Wing pictures of P) w1118, Q-W) expression of whole miR-309-6 cluster, miR-309-5 partial cluster, miR-6-1/2/3 partial cluster, or individual microRNAs in *patched* expressing cells. Abbreviations: Fe: femur; Ti: tibia; Ta: tarsus; Arrowhead indicates shortened tarsal segments; star indicates the loss of anterior wing vein.

To dissect if select microRNAs in the cluster were responsible for the altered leg phenotypes we generated homozygous UAS lines with subsets of members of the cluster; -*miR-309/3/286/4/5* (UAS-*miR-309-5*), UAS-*miR-286/4/5* and UAS-*miR-6-1/6-2/6-3*. Also, UAS lines were generated that expressed individual cluster members; UAS-*miR-309*, UAS-*miR-3* and UAS-*miR-286.* All lines were crossed with GAL4-*Dll* or GAL4-*ptc* drivers. Surprisingly, *Dll* or *ptc-*driven overexpression of either *miR-309-5* or *miR-6-1/6-2/6-3* (Fig. 3D, E, J and O, Supplementary information S4) recapitulated the phenotype created by overexpression of the entire cluster, while all other combinations showed normal leg phenotypes (Fig. 3 F-I and K-N). The number of tarsal segments were counted for different crosses, and only *Dll* or *ptc* driven overexpression of either *miR-309-5* or *miR-6-1/6-2/6-3* showed reduced tarsal segment numbers (Supplementary information S5). In addition to the aberrant tarsal segments, another phenotype of loss of the anterior wing cross vein was also observed in GAL4-*ptc*>*miR-309-6*, GAL4-*ptc*>*miR-309-5* and GAL4-*ptc*>*miR-6-1/6-2/6-3* flies (Fig. 3Q, R and W), but not with any of the other microRNA UAS lines (Fig. 3S-V). These data showed that the upregulation of either *miR-309-5* or *miR-6-1/6-2/6-3* partial clusters could cause the loss of tarsal segments and the anterior cross vein in a similar fashion to overexpression of the entire cluster.

UAS-microRNA-sponge lines that act as competitive inhibitors of the individual microRNAs including *miR-309, miR-3, miR-286, miR-4, miR-5* and *miR-6-1/6-2/6-3* were then crossed with GAL4-*Dll* and GAL4-*ptc*. None generated the leg or wing phenotypes, suggesting that the loss of any individual miRNA was insufficient to affect the development of leg and wing (Supplementary data S7). As loss-of-function of the whole cluster has been demonstrated to result in partial larval lethality and the survived adults possess normal leg phenotype (Bushati et al 2008; Chen et al 2014, this study); these results indicated that potential compensatory effects of other microRNA families might involve.

To understand which target genes are regulated by this microRNA cluster, total RNA was extracted from the leg discs of third instar larvae of GAL4-*ptc*>*miR-309-6* and GAL4-*ptc* (control), and subjected to Illumina Hi-Seq2500 sequencing. Third instar larvae were chosen as this is the developmental stage in which leg tarsal segments differentiate (Kojima 2004). Differentially expressed genes are shown in Supplementary data S8. Expression levels of CG32264 and CG10420 were similar in both the GAL4-*ptc*>*miR-309-6* and control, further reducing the possibility that the phenotypic change was caused by any effect on or of these genes.

Many of the genes involved in *Drosophila* leg development are known, and many of these were down-regulated in our transcriptome data (Supplementary data S8). Quantitative PCR was carried out to validate the gene expression changes in L3 leg discs of GAL4-*ptc*>*miR-309-6*, GAL4-*ptc*>*miR-309-5* and GAL4-*ptc*>*miR-6-1/6-2/6-3* (Fig. 4A-G). Our data showed that several leg patterning genes (such as *zfh-2*, *Sp1*, *Egfr*, *dysf*) were significantly down-regulated in mutant leg discs, suggesting repression of leg developmental genes by components of this microRNA cluster. UAS-RNAi lines of these down-regulated genes were crossed to GAL4-*ptc* and GAL4-*Dll*, and we found that *ptc*>*zfh-2*-RNAi, *Dll*>*zfh-2*-RNAi, *Dll*>*Sp1*-RNAi, *Dll*>*dysf*-RNAi, *ptc*>*Egfr*-RNAi, *ptc*>*dpp*-RNAi and *Dll*>*dpp*-RNAi resulted in tarsal segment deformities including loss of segment, joint boundaries and claws (Fig. 4H-N). *zfh-2* is a zinc finger homeodomain-2 transcription factor known to be involved in proximal-distal patterning of appendages (Guarner et al 2014), while the transcription factors *Sp1* and *dysf* are regulators of appendage growth and tarsal joint formation in insects (Córdoba and Estella 2014; Córdoba et al 2016). *Egfr* and *dpp* are also well known to be vital for leg patterning (Galindo et al 2002). These data indicate that the miR-309-5 and miR-6-1/2/3 sub-clusters target similar ‘leg development’ genes as the whole miR-309-6 cluster.

**Fig. 4.**
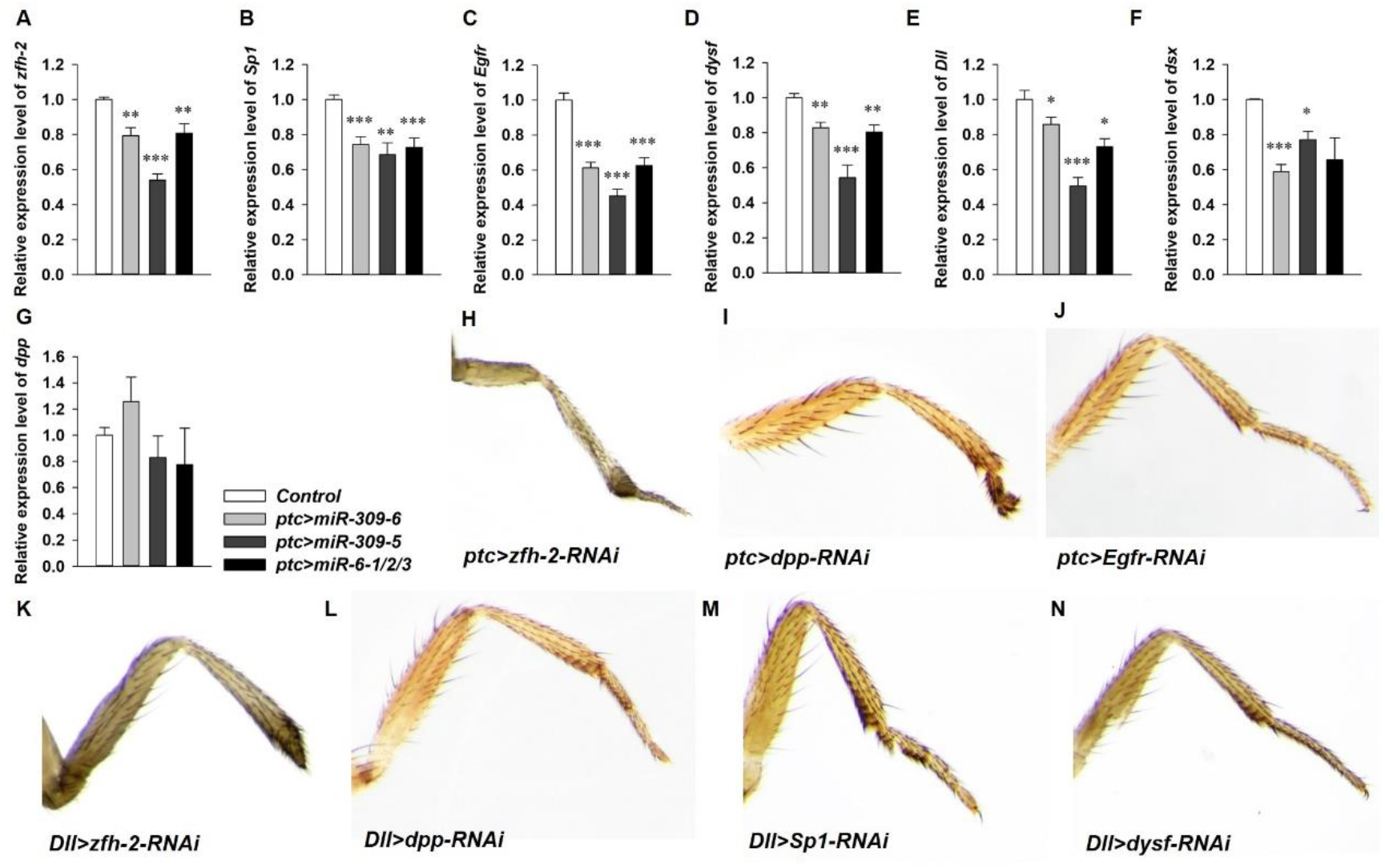
Expression of whole miR-309-6 cluster, or miR-309-5 and miR-6-1/2/3 sub-clusters regulates similar gene regulatory networks of leg development. A-G) Relative expression levels of *zfh-2*, *Sp1*, *Egfr*, *dysf*, *Dll, dsx* and *dpp* in leg discs of *ptc>miR-309-6*, *ptc>miR-309-5* and *ptc>miR-6-1/2/3*. Values represent mean ± S.E.M, * *p*<0.05, ** *p*<0.005 and *** *p*<0.001. H-J) Aberrant tarsal leg phenotypes created by *ptc* GAL4 driven expression of *zfh-2*-RNAi, *dpp*-RNAi and *Egfr*-RNAi. K-N) Tarsal leg deformities created by *Dll* GAL4-driven expression of *zfh-2*-RNAi, *dpp*-RNAi, *Sp1*-RNAi and *dysf*-RNAi.

To determine if there are other genes affecting leg development, two sets of differentially expressed genes were screened for further analyses, including 1) genes with significant expression change between controls and overexpression experiments, and 2) genes not expressed in the microRNA overexpression experiments but which are highly expressed in the controls. Twenty-four genes were identified as differentially expressed including *Arc1, Ag5r, Ag5r2, CG5084, CG5506, CG6933, CG7017, CG7252, CG7714, CG14300, CG17826, Eig71Eb, Hsp68, Hsp70Bb, Hsp70Bc, Mtk, Muc96D, Peritrophin-15a, Sgs3, Sgs5, stv, Obp99a, obst-I*, and *w* (Supplementary data S8). Genes that were absent or down-regulated in the GAL4-*ptc*>*miR-309-6* compared to controls were further tested by generating GAL4-*ptc* or *Dll*>UAS-RNAi lines for each gene, to check whether a short leg phenotype was observed. Similarly, GAL4-*ptc* or *Dll*>UAS-lines were generated for each gene that was upregulated in GAL4-*ptc>miR-309-6.* None of these individual manipulations were found to cause shortening of the leg or loss of tarsal segments (Supplementary data S9).

To further explore the genes being controlled by individual members of this miR-309-6 cluster, we transfected different combinations of the cluster microRNAs into *Drosophila* S2 cells and sequenced the transcriptomes. There were 178 differentially expressed genes in total when comparing to the controls (Fig. 5). Among these genes, 113 genes (~63.5%) and 65 genes (~36.5%) are commonly or uniquely regulated by microRNA cluster members, respectively (Supplementary data S10). Gene ontology (GO) enrichment analysis was carried out between the gene lists resulting from transfection of the whole cluster versus the younger members of the cluster (miR-6-1/2/3). There is no clear difference between the processes targeted by the whole cluster relative to the miR-6-1/2/3 sub-cluster, even in the ‘unique’ target category (Supplementary data S11).

**Fig. 5.**
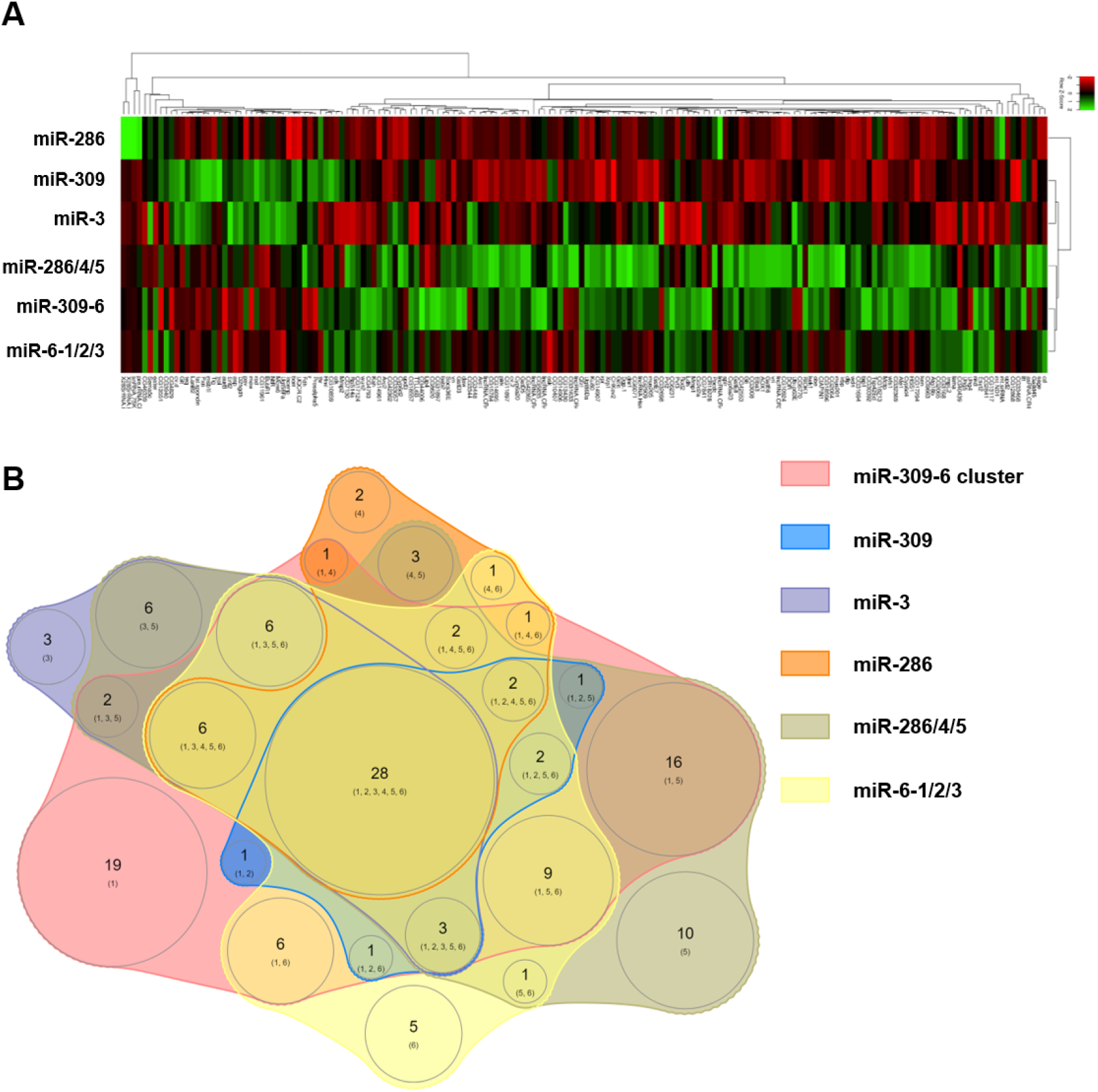
Cell transfection assays reveal common and unique targets of microRNAs in the miR-309-6 cluster. A) Transcriptome analyses of differential gene expression after transfection of different combinations of microRNAs from the cluster into *Drosophila* S2 cells. B) Venn-diagram showing the genes commonly or uniquely regulated by different combinations of microRNAs in the cluster. Numbers shown within the circles represent the genes being regulated by the relevant microRNA cluster members. Details of the gene list and their expression compared to the controls are listed in Supplementary data S10.

A question that has been frequently asked within the field is whether it is crucial and important for protein coding genes to be regulated by microRNAs. In some views, given that microRNAs can theoretically bind to hundreds of transcripts (e.g. Bartel 2009; Betel et al 2010; Reczko et al 2012), it has been proposed that the effect of microRNAs on targets would be weak and biologically irrelevant. In other views, based on the fact of sequence and target conservation and that some microRNAs have been found to strongly repress targets, which can result in phenotypic changes, it has also been argued that microRNA-protein-coding gene interactions are biologically significant. It is hypothesized that many microRNAs function to repress target transcription noise and stabilize gene regulatory networks, or can be important in evolution via such processes as microRNA arm switching (e.g. Flynt and Lai 2008; Pinzon et al 2017; Zhao et al 2017; Hornstein and Shomron 2006; Peterson et al 2009; Wu et al 2009; Ebert and Sharp 2012; Posadas and Carthew 2014; Marco et al 2010; Griffiths-Jones et al 2011; Hui et al 2013). In the *in vitro* and *in vivo* data provided here, expression of a polycistronic transcript containing eight microRNAs organised in a genomic cluster (miR-309-6) can result in phenotypic and gene expression changes when the microRNA expression is altered (Fig. 5 and 6A). Together with the fact that individual microRNAs alone cannot recapitulate the phenotypes, we conclude that action of the microRNA cluster as a combinatorial entity is important in gene regulation, and that future analyses should focus on the cluster and sub-cluster levels rather than on individual microRNAs alone.

**Fig. 6.**
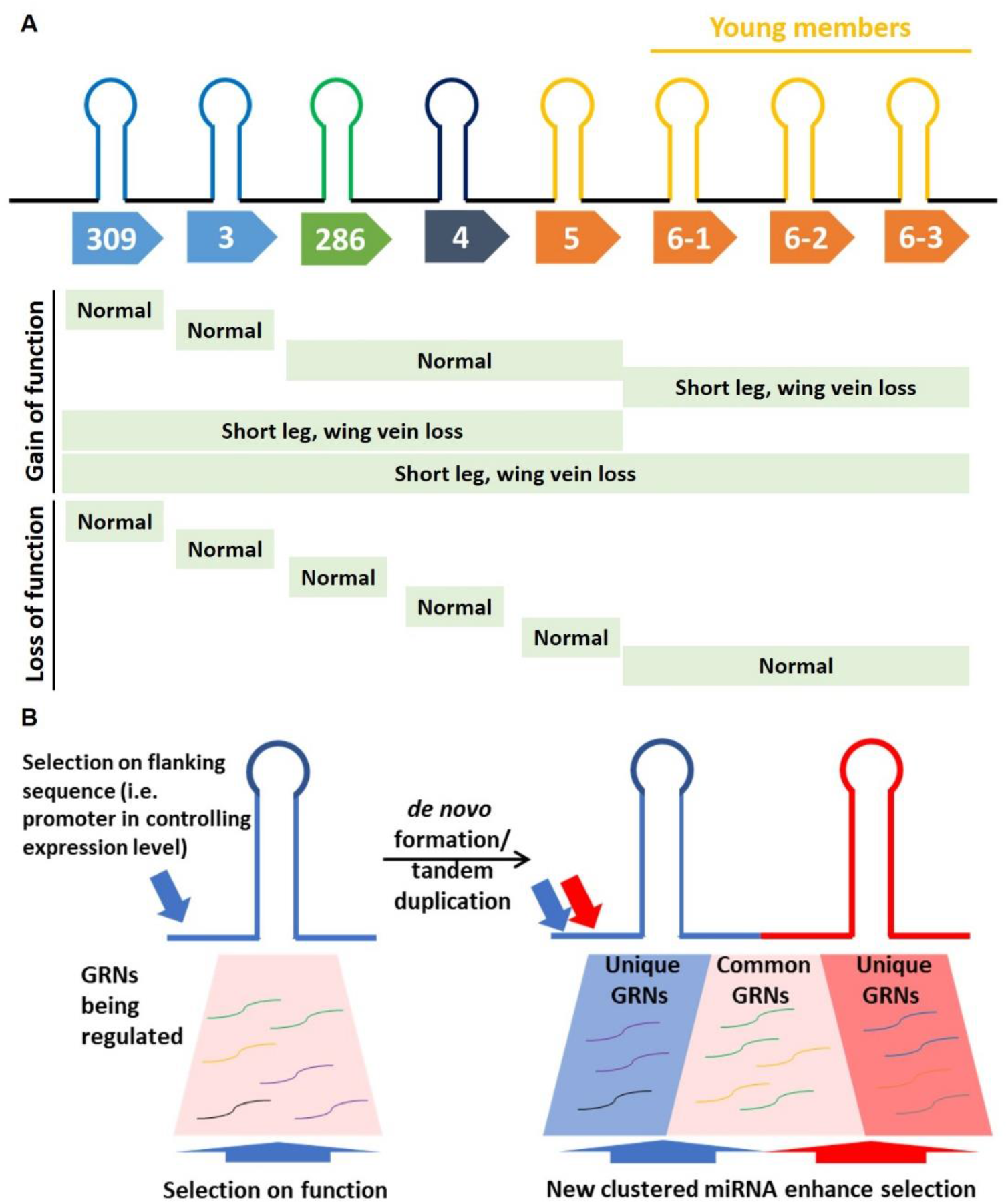
Varying selection forces acting via new and old microRNAs in a genomic cluster. A) Summary of the phenotypic results obtained in the gain-of-function and loss-of-function experiments of miR-309-6 cluster microRNAs in adult *Drosophila melanogaster*. Green colour depicts the GAL4/UAS mutants. B) During the formation of new microRNAs in a genomic cluster, selection forces are reinforced and potentially extended via the extension of the range of unique targets. For details, please refer to main text. The same colour denotes mRNAs involved in the same gene regulatory network (GRN). The arrows represent the selection forces acting on both the promoter sequence and microRNA sequences in an integrated fashion.

Another consideration is what the functional consequences of microRNAs being clustered are. Recently, it was shown that many mammal microRNA clusters may have multiple starting, end and processing sites, which not always transcribing all encoded microRNAs (Chang et al 2015; de Rie et al 2017). In *D. melanogaster*, miR-317/277/34 cluster have different primary microRNA isoforms (Zhou et al 2018). The miR-309-6 cluster was reported to be transcribed as a single transcript (Biemar et al 2005), which was supported by the Flybase RAMPAGE and ModENCODE CAGE data (Supplementary data S13). In addition, our 5’RACE and RT-PCR also validated the presence of the primary transcript of miR-309-6 cluster (Supplementary data S13). The conventional view focuses on the functional significance of shared targets by all microRNA members in a cluster, because with the regulation via a shared promoter, these microRNAs need to regulate their targets in the same cell at the same time. Hence, there is selection on the promoter of a microRNA cluster as members of the microRNA cluster must cooperatively function together (see introduction, Fig 6B). Our data support this view. For instance, if mutation occurred in the promoter sequence resulting in overexpression of this polycistronic transcript, this would then lead to phenotypes such as the shortening of legs. Such phenotypic changes would then potentially alter the organism’s fitness and be subjected to selection. This then can be viewed as an evolutionary constraint on the cluster promoter, with selective pressures on the promoter interacting with selective pressures on the evolution of the microRNA sequences themselves and the consequent evolution of the target affinities.

In addition to selection on the temporal and spatial aspects of microRNA cluster expression, levels of microRNA expression also appear to be functionally important and hence subject to selection. For example, when expressing only miR-309, miR-3, or miR-286-4-5, no phenotypic effects were observed, while the summation of expressing all of them (miR-309-5) resulted in phenotypic changes (Fig. 6A). These results suggest that as cluster composition evolves then selection on the promoter will also change, as ‘tuning’ of expression levels will likely be required in conjunction with changes to cluster membership (Fig. 6B). Given that expression of the younger members, miR-6-1/2/3, also results in similar phenotypes (Fig. 6A), it is likely that new microRNA members when arising in the microRNA cluster (via *de novo* formation or tandem duplication), can also enhance the selective pressures acting on the microRNA cluster. Another possibility is that individual microRNAs of a microRNA cluster can only target the leg patterning genes weakly, and a phenotype can only result when multiple leg genes are being targeted by multiple microRNAs in a cumulative manner. Plasticity-first evolution has been proposed as a predominant mechanism in nature (Levis and Pfennig 2016), and microRNAs have been postulated as a “missing link” in this process, by providing fine-tuning of expression networks and facilitating adaptation (Voskarides 2016). The evolution and functions of a microRNA cluster will then be a balance of sequence mutations on its promoter that control the spatiotemporal aspects and levels of cluster expression, and the functions of target genes either commonly or uniquely regulated by microRNAs inside the cluster. MicroRNA clusters must thus be viewed as integrated composites with both regulation and target affinities co-evolving in a concerted fashion.

## Methods

### Genome-wide target prediction

Mature miRNA sequences were retrieved from the public repository for published microRNA sequences at the miRBase database (http://www.mirbase.org). Eukaryotic 3’ untranslated region (UTR) sequences were retrieved from the public repository for published mRNA sequences at FlyBase (ftp://ftp.flybase.net/releases/FB2018_01/). All mature miRNAs were then used to predict targets in their respective genomes using the miRanda algorithm (Enright et al 2003) with parameters (i.e. -sc S Set score threshold to S 140 (from 140 to 772.00); -en -E Set energy threshold to -E kcal/mol (from −78.37 to −5.28); and -strict Demand strict 5’ seed pairing). For *D. melanogaster*, target prediction was also performed by Targetscanfly (Agarwal et al 2018).

### Fly culture, mutant construction, and insertion site checking

To prepare the overexpression constructs of *D. melanogaster* microRNA cluster miR-309-6 and miR-309-5, the corresponding stem-loop with flanking sequences was amplified and cloned into the GAL4-inducible vector pUAST (primer information is provided in Supplementary information S12). Constructs were sequenced prior to injection into *D. melanogaster w^1118^* embryos. Flies were screened and crossed to generate stable homozygous transformants. Insertion sites of the *UAS-miR-309-6* and *UAS-miR-309-5* cluster transgene were checked with Splinkerette PCR (Potter and Luo 2010). Various GAL4 drivers, UAS-gene and UAS-RNAi lines were obtained from the Bloomington *Drosophila* Stock Center (BDSC) (Supplementary information S6). The miR-309-6 whole cluster deletion line was ordered from BDSC (#58922, Chen et al 2014). UAS-CG32264-RNAi and UAS-CG10420-RNAi were donated by the Transgenic RNAi Project. UAS-miR-6-1/6-2/6-3, UAS-miR-286, UAS-miR-309 and UAS-miR-286/4/5 were donated by Stephen Cohen and Eric Lai, and UAS-miR-3 was obtained from the Zurich ORFeome Project. UAS-microRNA-sponge lines including miR-3, miR-4, miR-5, miR-6-1/2/3, miR-286 and miR-309 were donated by David Van Vactor. All flies were maintained on standard yeast-cornmeal-agar medium at 25°C. Males and virgin females from each fly line were randomly collected for crossings. For each crossing of GAL4 and UAS fly lines, three random males and three random virgin females were used, and reciprocal crosses were carried out. At least three separate crossings were performed for each GAL4 and UAS pair.

### MicroRNA expressing vector construction and cell transfection

MicroRNAs were amplified from *D. melanogaster* (primer information shown in Supplementary information S12). Amplicons were cloned into pAC5.1 vector (Invitrogen). All constructs were sequenced to confirm their identities. *Drosophila* S2 cells (DRSC) were kept at 23°C in Schneider *Drosophila* medium (Life Technologies) with 10% (v/v) heat-inactivated fetal bovine serum (Gibco, Life Technologies) and 1:100 Penicillin-Streptomycin (Gibco, Life Technologies). The pAC5.1-miRNA (300 ng) was transfected into *Drosophila* S2 cells using Effectene (Qiagen) per the manufactures’ instructions. RNA was isolated at 48 h post-transfection.

### Transcriptome and small RNA sequencing

RNA was extracted from leg discs of pupariating 3^rd^ instar larvae of GAL4-*ptc*>*miR-309-6*, GAL4-*ptc*>*miR-309-5*, GAL4-*ptc*>*miR-6-1/6-2/6-3* and GAL4-*ptc* (control) lines, S2 cells expressing different combination of microRNAs in the cluster and S2 (control) cells. Wildtype leg disc RNA was processed by BGI for HiSeq Small RNA library construction and 50 bp single-end (SE) sequencing. The expressed microRNAs were quantified by the “quantifier.pl” module of the mirDeep2 package with parameters “-g 0 -e 0 -f 0”, and the clean reads were aligned to hairpin sequence from miRBase (release 22) by bowtie with parameters “-l 18 -v 0 -a --best --norc --strata”. Transcriptome libraries were constructed using the TruSeq stranded mRNA LT sample prep kit, and sequenced on an Illumina HiSeq2500 platform (BGI Hong Kong). Raw reads were filtered using Trimmomatic and mapped to Flybase v.6.14 using Cufflinks (Trapnell et al 2012). Differential gene expression was evaluated using Cuffdiff.

### Taqman microRNA assays and real-time PCR

Expression of microRNAs was measured via Taqman microRNA assays (Applied Biosystems™) following the manufactures’ instructions. For detection of differential gene expression, RNAs from the respective crosses were reverse-transcribed into cDNA using the iScript™ cDNA synthesis Kit (BioRad). Real-time PCR was conducted in three biological replicates using the CFX96 Touch™ Real-Time PCR Detection System (BioRad), with a programme of denaturation at 95°C for 3 min followed by 40 cycles of 95°C/ 10s, 55°C/ 10s and 72°C/ 15s. PCRs were run with half iTaq™ Universal SYBR^®^ Green Supermix (BioRad) and 0.2 μM of each primer pair (primer information is listed in Supplementary information S12).

## Supplementary data / information

S1. Sequence alignment of microRNAs in miR-309-6 cluster.

S2. Target gene prediction of different microRNAs in microRNA-309-6 cluster of different *Drosophila* genomes.

S3. Expression of miR-309-6 cluster microRNAs in *D. melanogaster*.

S4. Tarsal segments of *Dll* and *ptc* GAL4-driven expression of miR-309-6, miR-309-5 and miR-6-1/2/3.

S5. Tarsal segment numbers of *Dll* and *ptc* GAL4-driven expression of miR-309-6, miR-309-5 and miR-6-1/2/3 mutants.

S6. UAS-gene and UAS-RNAi fly lines used in this study.

S7. Leg and wing pictures of loss-of-function of individual microRNAs from the miR-309-6 cluster in either the *Distal-less* (A-F) or *patched* (G-R) expressing cells.

S8. Differential expression of genes with over 2-fold changes and leg patterning related genes between GAL4-ptc>miR-309-6 (mutant) and control (GAL4-ptc) leg discs.

S9. Phenotypes of gain-and loss-of-function of selected differentially expressed genes.

S10. List of genes being regulated by microRNA cluster members *in vitro*.

S11. Gene Ontology (GO) enrichment analysis

S12. Primer information used in this study.

S13. Evidences showing the polycistronic transcript of miR-309-6 cluster.

## Author Contributions

JHLH conceived and supervised the study. ZQ, WCY, HYY, CWCY, WN, AYPW, IHTL, JHLH carried out the experimental works. NWYW cloned the cluster into pUAST vector. FKMC obtained homozygous red-eye lines after pUAST vector injection by BestGene. ZQ, WCY, HYY, CWCW, WN, AYPW, IHTL, FKMC, TFC, KFL, SZ, KHC, DEKF, WGB, JHLH wrote the manuscript.

## Acknowledgements

The authors thank KB Wong and HM Lam for discussion. This research was supported by the Hong Kong Research Grant Council GRF Grant (14103516), The Chinese University of Hong Kong Direct Grant (4053248), and TUYF Charitable Trust (6903957) (JHLH).

